# Meningeal origins and dynamics of perivascular fibroblast development on the mouse cerebral vasculature

**DOI:** 10.1101/2023.03.23.533982

**Authors:** Hannah E. Jones, Vanessa Coelho-Santos, Stephanie K. Bonney, Kelsey A. Abrams, Andy Y. Shih, Julie A. Siegenthaler

## Abstract

Perivascular fibroblasts (PVFs) are a fibroblast-like cell type that reside on large-diameter blood vessels in the adult meninges and central nervous system (CNS). PVFs drive fibrosis following injury but their homeostatic functions are not well detailed. In mice, PVFs were previously shown to be absent from most brain regions at birth and are only detected postnatally within the cerebral cortex. However, the origin, timing, and cellular mechanisms of PVF development are not known. We used *Col1a1-GFP* and *Col1a2-CreERT* transgenic mice to track PVF developmental timing and progression in postnatal mice. Using a combination of lineage tracing and *in vivo* imaging we show that brain PVFs originate from the meninges and are first seen on parenchymal cerebrovasculature at postnatal day (P)5. After P5, PVF coverage of the cerebrovasculature rapidly expands via mechanisms of local cell proliferation and migration from the meninges, reaching adult levels at P14. Finally, we show that PVFs and perivascular macrophages (PVMs) develop concurrently along postnatal cerebral blood vessels, where the location and depth of PVMs and PVFs highly correlate. These findings provide the first complete timeline for PVF development in the brain, enabling future work into how PVF development is coordinated with cell types and structures in and around the perivascular spaces to support normal CNS vascular function.

**Summary:** Brain perivascular fibroblasts migrate from their origin in the meninges and proliferate locally to fully cover penetrating vessels during postnatal mouse development.

## INTRODUCTION

The central nervous system (CNS) requires continuous delivery of blood containing oxygen and nutrients. This is ensured by a robust vascular network made up of many different cell types—endothelial cells, pericytes, vascular smooth muscle cells (vSMC), astrocytes, immune cells, and perivascular fibroblasts (PVFs)—which come together to form the specialized neurovascular unit. The endothelium, pericytes, vSMCs, and astrocytes have all been well studied in the context of development of the neurovascular unit and their developmental origins are known (Coelho-Santos and Shih, 2020; Paredes et al., 2018). In contrast, much less is known about how perivascular fibroblasts (PVFs) develop, integrate, and function in the neurovasculature.

PVFs are a fibroblast-like cell type located on the outside of the vSMC layer of large-diameter arterioles and venules, but not capillaries, in the adult mouse brain (Bonney et al., 2022; Soderblom et al., 2013; Vanlandewijck et al., 2018) and in adult human brain (Garcia et al., 2022; Yang et al., 2022). PVFs express collagen-1 and are distinct from other mural cells (pericytes and vSMCs) in their expression of both platelet derived growth factor (PDGF) receptors α and β and absence of mural cell markers including desmin and NG2 (Fernández-Klett et al., 2013; Kelly et al., 2016; Riew et al., 2020; Soderblom et al., 2013; Vanlandewijck et al., 2018). Given their position within the neurovascular unit, PVFs are uniquely positioned to provide support to vascular structures and cell types, potentially via secretion or modification of extracellular matrix proteins like type I and III collagens (Hannocks et al., 2018; Vanlandewijck et al., 2018; Yang et al., 2022). Most research on PVFs has focused on their robust CNS injury and disease response. Our lab and others have shown that, following acute CNS injuries like stroke and traumatic brain injury, PVFs detach from the vasculature, proliferate, and contribute to fibrotic scar formation (Fernández-Klett et al., 2013; Kelly et al., 2016; Soderblom et al., 2013). Additionally, there is evidence that PVFs contribute to pathologies seen in neurodegeneration and neuroinflammation (Dorrier et al., 2021; Månberg et al., 2021). Despite known roles of PVFs in CNS injury and disease, there have been few insights into the cellular mechanisms of their development and how this relates to other neurovascular cell types. Previous work from our lab showed that PVFs are absent from the brain vasculature at birth and only appear postnatally (Kelly et al., 2016), many days after other cell types assemble a functional neurovascular unit. An important step towards understanding the homeostatic and disease-response functions of PVFs is to identify the cellular mechanisms that underlie PVF development, and how PVF development relates to other aspects of neurovascular function.

The *Collagen1a1(Col1a1)-GFP* and *Collagen1a2(Col1a2)-CreERT* mouse lines are established tools for labeling and identifying PVFs in the mouse brain, as they are expressed in all CNS fibroblasts (Bonney et al., 2022; Dorrier et al., 2021; Soderblom et al., 2013). Here, we use these tools to investigate PVF development in the cerebral cortex during postnatal development. We show that *Col1a1-GFP+* PVFs appear on leptomeningeal vessels embryonically, but do not start to cover vessels in the cerebral cortex until postnatal day (P)5 and reach full coverage by P14. We also show that timing of brain PVF appearance aligns with the timing of another CNS perivascular cell type, perivascular border associated macrophages (PVMs), with PVF and PVM coverage of cerebral vessels occurring in tandem. Lineage tracing using the *Col1a2-CreERT* allele along with *tdTomato-flox* and *Brainbow-flox* reporters show that PVFs in the brain originate from the meninges and have distinct clonal boundaries on cerebral vessels, suggesting specific temporal and regional contributions of PVFs to vessel coverage. Further, *in vivo* two-photon imaging and proliferation assays reveal that both migration and proliferation drive PVF coverage of vessels. From these data emerge the first comprehensive timeline of PVF development in the postnatal cerebral cortex, enabling a more complete model of postnatal cerebrovascular development and insight into how PVF development may be coordinated with other perivascular cell types and structures.

## RESULTS

### PVFs emerge on cerebral vessels between P5 and P14 in the mouse cerebral cortex

To create a timeline of PVF emergence in the postnatal mouse cerebral cortex, we used the *Col1a1-GFP* transgenic mouse line, in which all CNS fibroblasts including PVFs are labeled, along with CUBIC tissue clearing of 2-mm brain slices to observe PVF locations on vessels at different timepoints. In the adult mouse brain, PVFs are present on large-diameter vessels across all brain regions (Bonney et al., 2022; Kelly et al., 2016). We analyzed non-capillary penetrating vessels in the cerebral cortex, as it has been established that PVFs localize to penetrating arterioles and ascending venules but not capillaries in mice (Bonney et al., 2022). At postnatal day (P)5 *Col1a1-GFP+* fibroblasts are abundant in the meninges and along the hippocampal vasculature but are rare in the cortex (**Fig. 1A**). At this age, approximately 15% of all penetrating vessels in the cortex have some amount of coverage by *Col1a1-GFP+* PVFs (**Fig. S1A**) and the PVFs present are located close to the meningeal surface, with a mean depth of only about 11µm into the cortex (**Fig. 1B**). Occasionally at P5, individual PVFs are observed proximal to the meninges, initiating their coverage on vessels (**Fig. 1A**, P5, arrowhead). At P7 and P10, GFP+ PVFs continue emerging and covering vessels (**Fig. 1A**, P7 & P10). The percent of penetrating vessels with some amount of PVF coverage at P7 and P10 increases significantly, from about 15% at P5 to 60% at P7 (p = 0.0136, p < 0.05 *), and to 99% at P10 (p = 0.006, p < 0.01 **) (**Fig. S1A**). By P14 GFP+ PVFs reach the terminus of vessels (**Fig. 1A**, P14 inset, arrowheads) and all cerebral penetrating vessels contain PVFs (**Fig. S1A**). In adults, PVFs are found on the main trunk of vessels and on arteriole branches up to the 3^rd^ order (Bonney et al., 2022). Interestingly, we observed that PVFs only cover vessel branches after covering the primary trunk of the vessel (**Fig. 1A**, P14 inset, arrow). These results are consistent with our prior work using Collagen-1 protein expression to observe appearance of fibroblasts between P0 and P21 in the mouse brain (Kelly et al., 2016).

**Figure 1:**
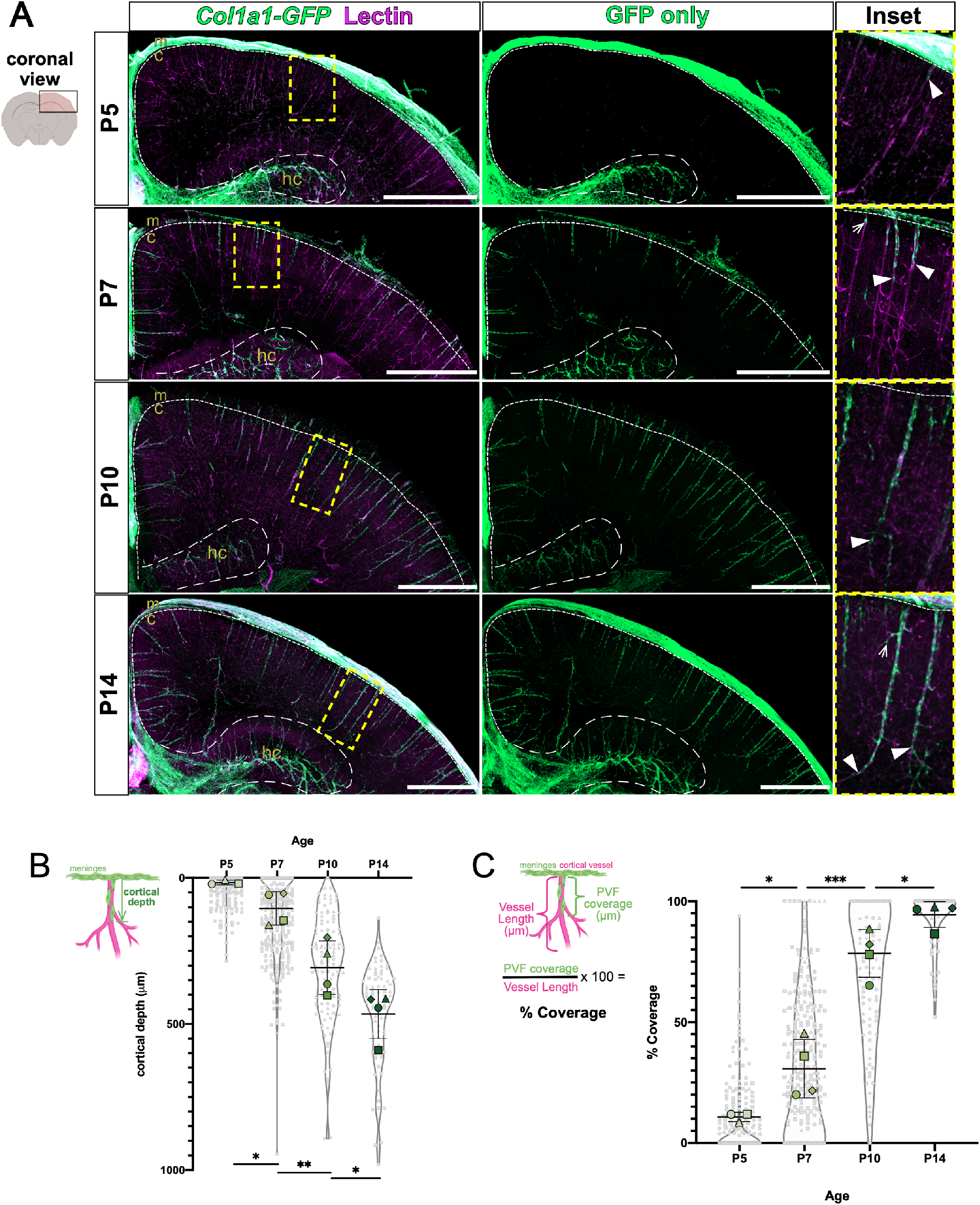
*Col1a1-GFP+* PVFs increase in depth in the cerebral cortex to cover large-diameter vessels between P5 and P14. **(A)** Coronal sections of CUBIC-cleared P5, P7, P10, and P14 *Col1a1-GFP* brains showing fibroblasts (GFP, green) and vessels (Lectin, magenta). In insets, arrowheads mark the position of the furthest-migrated GFP+ PVF on individual vessels at each timepoint. In P7 inset, arrow indicates PVF emerging on vessel and in P14 inset, arrow indicates PVF extending on to vessel branch. Note that decreased GFP fluorescence above the cortex in P7 and P10 images was due to meninges damage during dissection or sectioning. Meninges, **m**; cortex, **c**; hippocampus, **hc**. Scale bars 1mm. **(B-C)** Graphs depicting the **(B)** maximum depth of the furthest-migrated PVF on vessels measured from the meninges at ages P5, P7, P10, and P14 and **(C)** percent coverage of vessels at each age. For **B-C**, the means for each biological replicate (n = 3 – 4 for each age) is represented by large colored shapes and corresponding technical replicates (n = 15-255 vessels per animal) in small gray matching shapes, line and error bars show mean and SD. One-way ANOVA with multiple comparisons indicates statistically significant differences in cerebral depth and coverage between each timepoint (**B)** P5 vs. P7, p =0.0468; P7 vs. P10, p =0.0094; P10 vs. P14, p = 0.0434; **C)** P5 vs. P7, p =0.0393; P7 vs. P10 p =0.0009; P10 vs. P14, p = 0.0289; p < 0.05 *, p < 0.01 **, p < 0.001 ***).

During PVF emergence on vessels between P5 and P14, they appear deeper within the cortex at each timepoint. To investigate the dynamics of PVF appearance, we first measured the cortical depth of the furthest-traveled PVF on individual vessels from the meningeal surface at P5, P7, P10, and P14 (**Fig. 1B**). At P5, PVFs remain close to the meningeal surface, with a mean depth of about 11µm (**Fig. 1B**). PVF depth on vessels increases significantly between P5 to P7 (P7 mean = 99.3µm, p =0.0468, p < 0.05 *), and P7 to P10 (P10 mean = 300.4µm, p =0.0094, p < 0.01 **) (**Fig. 1B**). By P14, PVFs have reached a mean depth of about 457µm, this is significantly increased from P10 (p = 0.0434, p < 0.05 *) (**Fig. 1B**). We also conducted analysis of the extent of each vessel covered by PVFs at each timepoint by measuring the length of PVF coverage on individual vessels and dividing this measurement by the total vessel length to yield percent coverage (**Fig. 1C**). Consistent with our depth analysis, at P5, PVFs remain close to the surface of the cortex, with the mean vessel coverage at 3% (**Fig. 1C**). This increases significantly from P5 to P7, with the mean percent coverage increasing to 25% (p =0.0393, p < 0.05 *) (**Fig. 1C**). Between P7 and P10, PVFs significantly increase their coverage of vessels from 25% to 77% (p =0.0009, p < 0.001 ***) (**Fig. 1C**); this corresponds with the timespan in which there is the greatest increase in the depth of PVFs on vessels (**Fig. 1B**, P7 to P10). Between P10 and P14, PVF coverage again significantly increases from 77% to 94% (p = 0.0289, p < 0.05 *). We measured the density of PVFs within the covered regions on vessels and found no significant changes in cell density across timepoints (**Fig. S1B**), indicating PVFs on cerebral vessels increase their overall numbers to cover a greater length of vessel. Of note, did not distinguish between arterioles and venules in our analysis. Previous work on PVFs in adult mice show that venules tend to have less PVF coverage than arterioles (Bonney et al., 2022), this is likely reflected by data points of PVFs in **Fig. 1B-C** that have less than average depth or coverage. Taken together, these data show that PVF appearance between P5 and P14 corresponds with an increase in the depth of PVFs on vessels and an increase of coverage of cerebral vessels without any change in PVF density.

### Lineage tracing reveals a meningeal origin for cortical PVFs

Our analysis of postnatal *Col1a1-GFP+* brains shows that PVFs are absent from the brain prior to P5 but are present in the meninges residing directly above the cortical vessels they will eventually populate. This sets up the question; do brain PVFs originate from the meninges? To test this, we conducted lineage tracing using the inducible *Col1a2-CreERT2* transgenic mouse line that is expressed by fibroblasts in the CNS but not other perivascular cell types like vSMCs (Bonney et al., 2022; Dorrier et al., 2021; Zheng et al., 2002). We combined the *Col1a2-CreERT* line with an *Ai14(tdTomato)-flox* reporter to achieve fibroblast-specific expression of tdTomato upon introduction of tamoxifen. We injected *Col1a2-CreERT;tdTomato-flox* pups with a saturating dose of tamoxifen (e.g. a high enough dose to cause recombination in all fibroblasts) at P1, several days before any PVFs emerge on cerebral vessels, and collected brains for CUBIC-clearing and analysis at P3, P7 and P14 (**Fig. 2**). At P3, the only *tdTomato+* fibroblasts observed are in the meninges and around the hippocampal vasculature and are completely absent from vessels in the cortex (**Fig. 2**, P3 inset). This is consistent with what we would expect to see, given that in the *Col1a1-GFP* mouse line at P5, GFP+ fibroblasts are almost exclusively localized to the meninges and the hippocampal vasculature and have not yet entered the cortex (**Fig. 1A**). At P7, we begin to see *tdTomato+* PVFs appear on vessels in the cortex at depths similar to what we see in the *Col1a1-GFP* line at the same timepoint (**Fig. 2**, P7 inset, arrowheads), and by P14 we see *tdTomato+* PVFs extensively covering the cerebral vessels in addition to labeling of cells in the meninges, mirroring what we observe with the *Col1a1-GFP* line at P14 (**Fig. 2**, P14 inset, and **Fig. 1A**). Thus, PVFs in the brain developmentally originate from fibroblasts in the meninges.

**Figure 2:**
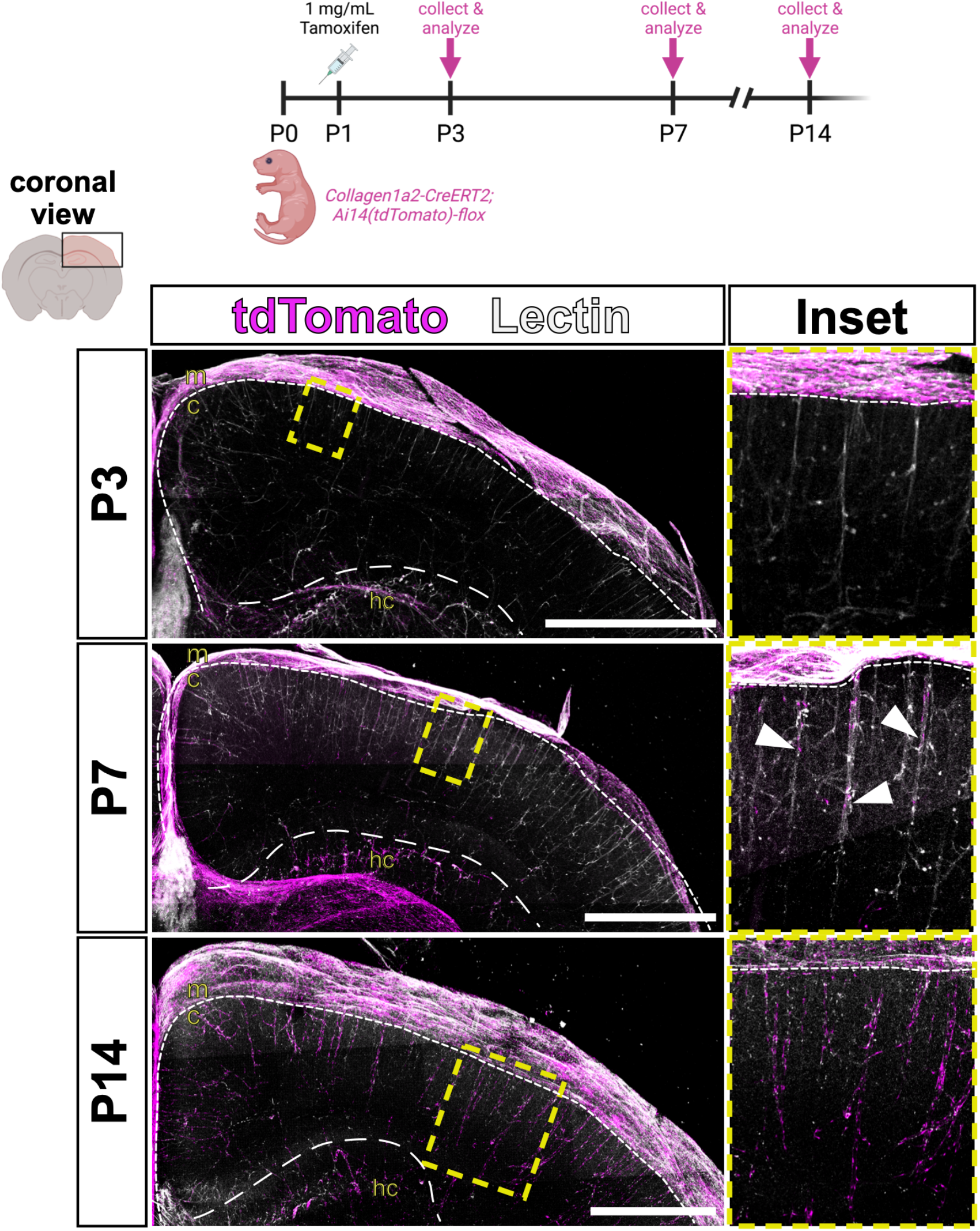
Parenchymal PVFs originate in the meninges. Lineage tracing of fibroblasts in coronal sections of CUBIC-cleared *Col1a2-CreERT2;Ai14-flox* brains showing vessels (Lectin, white) and fibroblasts recombined with tamoxifen induction at P1 (tdTomato, magenta) and followed at P3, P7, and P14. Arrowheads in P7 inset indicate the furthest-migrated *tdTomato+* PVFs on each vessel shown. Meninges, **m**; cortex, **c**; hippocampus, **hc**. Scale bars 1mm.

### PVFs first appear in the leptomeninges of embryonic and early postnatal mouse brains

*Col1a1-GFP+* PVFs do not appear on cerebral vessels until P5 but are abundant in the meninges at prior ages (**Fig. 1A, Fig. 2**). To better understand the origins of brain PVFs, we next investigated PVFs in the meninges prior to P5. The pial layer of the leptomeninges contains an extensive vascular network that gives rise to the penetrating vessels of the cortex (**Fig. 3A**). Our lab has pioneered techniques for flat-mount visualization of the leptomeninges (Jones et al., 2022), which allows us to observe and distinguish between non-perivascular and perivascular meningeal fibroblasts based on localization and morphology (**Fig. 3A**). We first sought to understand when PVFs first begin to appear in the leptomeninges; to investigate this we prepared flat-mounts of the leptomeninges at embryonic day (E)12 (**Fig. 3B**). At this time, the meninges are a meshwork of *Col1a1-GFP+* fibroblasts containing the perineural vascular plexus that surrounds the developing CNS but has not yet begun to specify into molecularly distinct meningeal layers (DeSisto et al., 2020). Even though the fibroblasts at E12 remain mostly undifferentiated, we observe meningeal PVFs with characteristic elongated morphology appearing along vessels at this timepoint (**Fig. 3B** inset, arrowheads). As meningeal layer-specific markers begin to emerge around E14-15 (DeSisto et al., 2020), PVFs continue to associate with meningeal vessels and do not express the mural cell marker Desmin (**Fig. 3C** image, arrowheads). Staining for protein expression of layer-specific markers S100A6 (pia-specific marker) and RALDH2 (arachnoid-specific marker) in E15 flat-mounts reveals that meningeal PVFs express both markers (**Fig. S2A-B**, arrowheads), although expression of both markers within the meningeal PVF population appears to be heterogeneous. We measured the density of *Col1a1-GFP+* PVFs on meningeal vessels at E14, E16, P0, and P2 and found that their density does not significantly increase from E14 through P0 (**Fig. 3D**). Together, these results suggest that during embryonic development, meningeal PVFs associate with the meningeal vasculature and reflect a morphologically and spatially distinct population from non-perivascular meningeal fibroblasts and other perivascular cell types, in particular mural cells.

**Figure 3:**
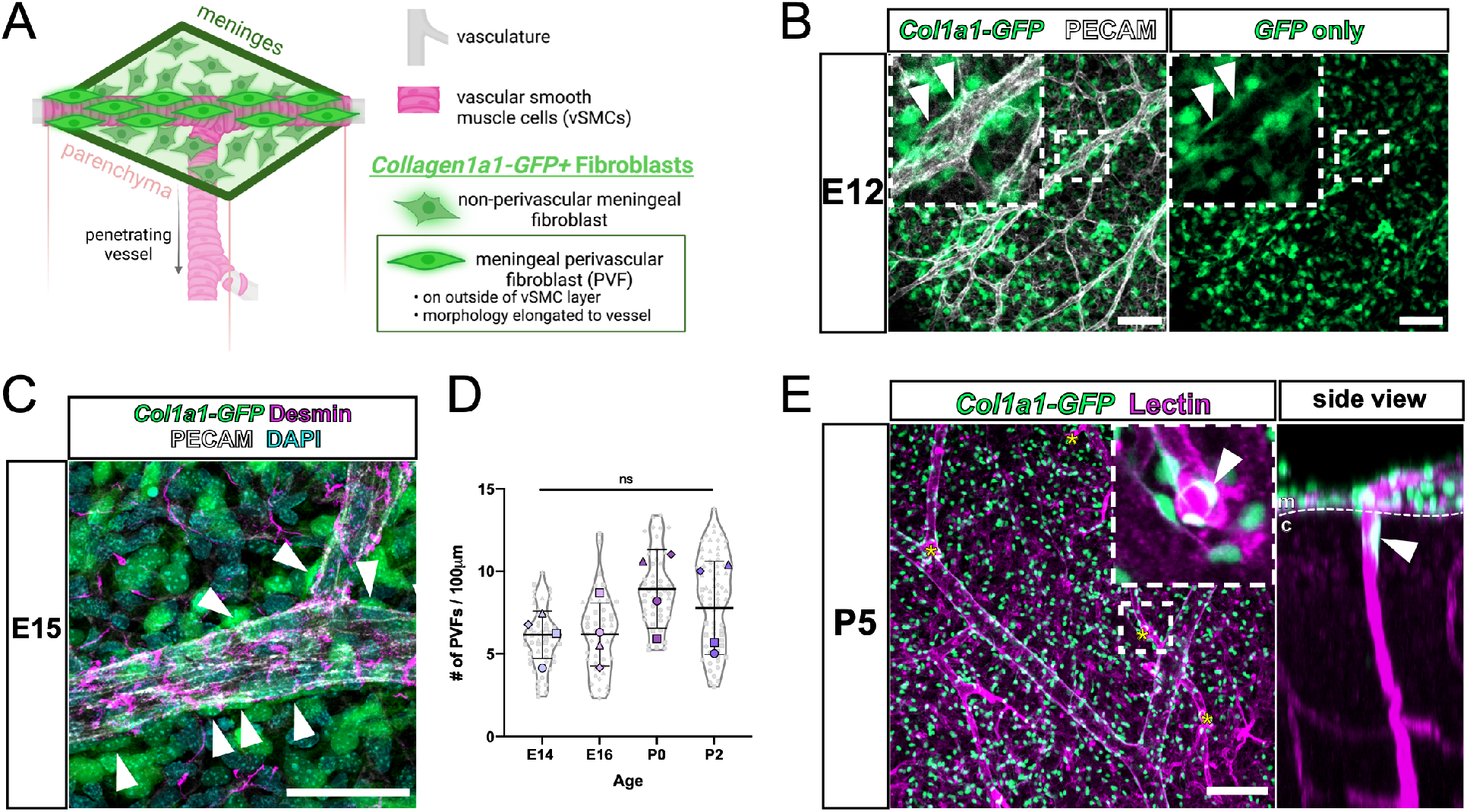
PVFs are present on cerebral leptomeningeal vessels embryonically prior to initiating coverage of parenchymal cerebral vessels at P5. **(A)** Model depicting location and characteristics of meningeal PVFs prior to their development on parenchymal vessels. **(B)** Flat-mount preparation of E12 meninges showing fibroblasts (GFP, green) and vasculature (PECAM, white). Inset and arrowheads mark meningeal PVFs displaying perivascular location and elongated morphology. Scale bars 100µm. **(C)** Flat-mount preparation of E15 meninges showing fibroblasts (GFP, green), vasculature (PECAM, white), pericytes/vSMCs (Desmin, magenta), and nuclei (DAPI, cyan). Arrowheads mark meningeal PVFs. Scale bar 50µm. **(D)** Graph depicts density (number of cells per 100µm of vessel length) of PVFs on meningeal vessels at E14, E16, P0, and P2, with the means of each biological replicate (n = 4 for each age) represented by large colored shapes and corresponding technical replicates (n = 5-28 vessels per animal) in small gray matching shapes, line and error bars show mean and SD. One-way ANOVA with multiple comparisons revealed no significant changes across timepoints. **(E)** Low-magnification scan of the meninges of a CUBIC-cleared P5 *Col1a1-GFP* cortex, showing fibroblasts (GFP, green) and vasculature (Lectin, magenta). Yellow asterisks mark vessels penetrating the cortex containing parenchymal PVFs located proximal to the meningeal surface (inset, arrowhead). Side view is a projection of the yz-plane of the inset showing the penetrating vessel and PVF (arrowhead). Meninges, **m**; cortex, **c**. Scale bar 100µm.

Blood vessels in the meninges are directly connected to the penetrating vessels of the cortex (**Fig. 3A**). Our data shows that up until P5, PVFs remain associated with vessels in the meninges. To investigate the localization of PVFs at P5, we visualized the meningeal vasculature and cerebral penetrating vessels concurrently using a whole-cortex clearing and imaging protocol (**Fig. 3E**). Here, we observe penetrating vessels branching from meningeal vessels (**Fig. 3E**, asterisks). Each of these penetrating vessels is wrapped with a *Col1a1-GFP+* PVF proximal to the meningeal surface (**Fig. 3E**, inset and side view, arrowheads). We referred to these PVFs seen on penetrating vessels proximal to the meningeal surface as “initiating PVFs”, as they are the first PVFs to appear on cerebral vessels and are initiating coverage of vessels at this timepoint. Combining information from our embryonic meningeal (**Fig. 3A-C**) and postnatal analyses (**Fig. 1**), P5 appears to be a critical timepoint in PVF development in which meningeal PVFs initiate coverage of cerebral vessels.

### PVFs and Perivascular Macrophages (PVMs) develop concurrently in the postnatal mouse brain

Recent work showed a subset of CNS border-associated macrophages called perivascular macrophages (PVMs), defined by expression of CD206 and location in perivascular spaces of large diameter vessels, arise from CD206+ leptomeningeal macrophages and appear postnatally (P5-P21) on vessels in the mouse brain (Karam et al., 2022; Masuda et al., 2022). Given the similar timing to the PVF development, we analyzed the dynamics of PVF and PVM progression onto individual cerebral vessels by applying similar analyses as conducted on PVFs in **Figure 1**.

First, we determined the relative location, depth, and density of CD206+ PVMs and *Col1a1-GFP+* PVFs on cerebral vessels at P6, P7, P8, and P9 (**Fig. 4**). At P6 and P7, we see both PVMs and PVFs localized to vessels (**Fig. 4A**), with most vessels containing both cell types (**Fig. 4B**; P6; Both = 81.82% ; P7, Both = 91.30%). Notably at P6 and P7, PVFs are observed slightly deeper along vessels compared to PVMs (**Fig. 4A**, P6 and P7, arrowheads) and our analysis shows most vessels at P6 and P7 timepoints have a PVF located deeper than a PVM (**Fig. 4C**; P6; PVF Deepest = 67.05%; P7, PVF Deepest = 66.67%; **Fig. S3A**). Further, a greater percentage of cerebral vessels had only a PVF at P6 compared to P7 (**Fig. 4B**; P6: PVF only = 15.91%, PVM only = 2.27%; P7: PVF only = 7.25% PVM only = 1.45%%). Like PVFs, PVMs follow a similar pattern with respect to their depth on vessels across timepoints, with PVMs significantly increasing their depth on vessels from P6 to P9/10 (p = 0.0097, p < 0.01 **) (**Fig. 4D**), however the average depth of PVF compared to PVMs on all cerebral vessels analyzed at P6, P7, P8 or P9-10 was not significantly different (**Fig. 4D**). By P9-10, all cerebral vessels analyzed have both PVFs and PVMs present (**Fig. 4B**) but PVFs were more often observed slightly deeper than PVMs (**Fig. 4C**). This data points to a model where PVFs initiate coverage of cerebral vessels ahead of PVMs but is quickly followed by PVM appearance.

**Figure 4:**
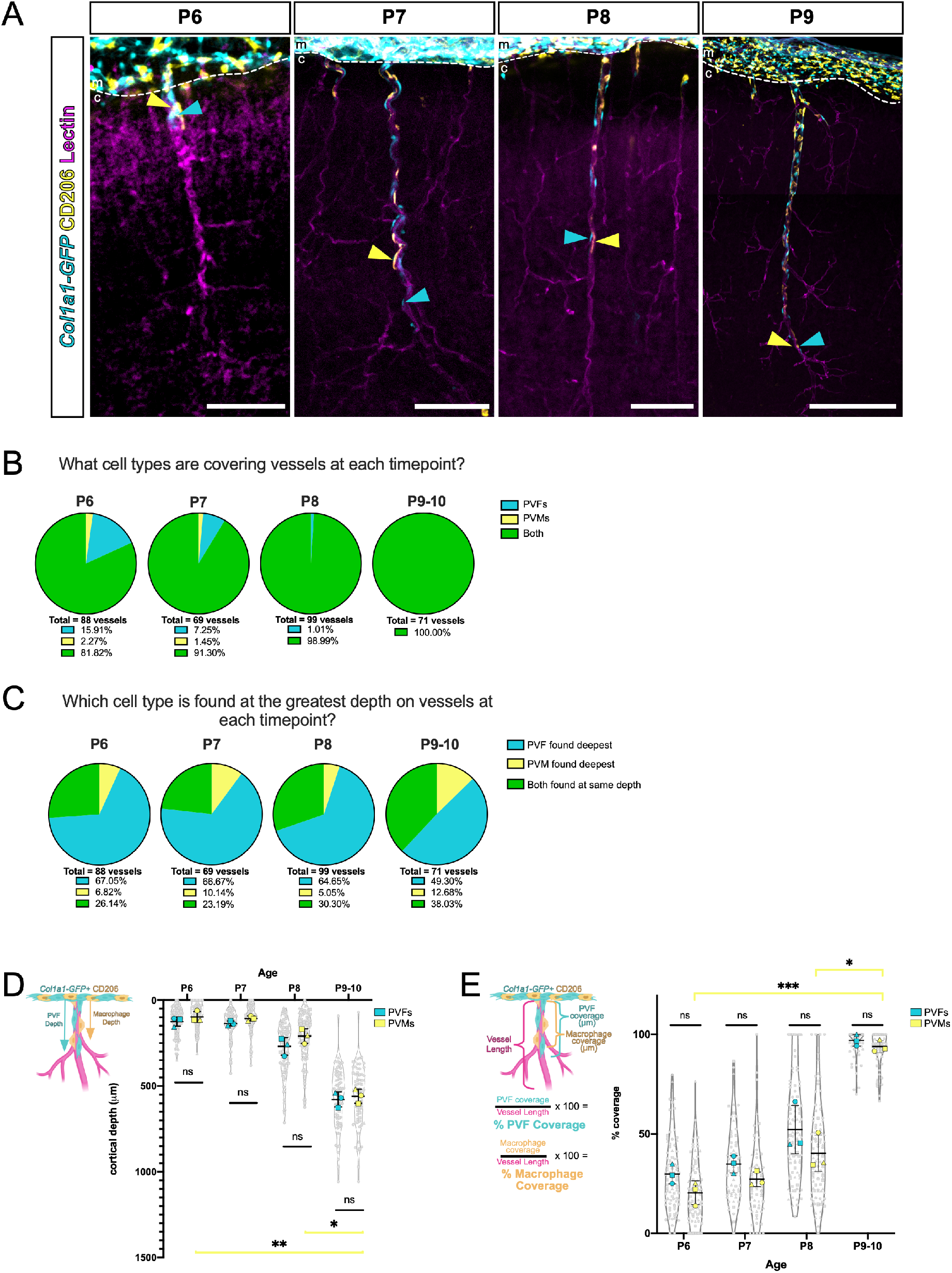
*Col1a1-GFP+* PVFs appear concurrently with CD206+ PVMs on cerebral vessels postnatally. **(A)** High-magnification images of individual vessels of CUBIC-cleared *Col1a1-GFP* brains at P6, P7, P8, and P9 showing fibroblasts (GFP, cyan) and perivascular macrophages (PVM) (CD206, yellow) on vessels (Lectin, magenta). In each panel, arrowheads mark the position of the furthest-migrated GFP+ PVF (cyan arrowhead) and CD206+ PVM (yellow arrowhead) on individual vessels at each timepoint. Meninges, **m**; cortex, **c**. Scale bars 100µm. **(B-C)** Pie-charts representing parts-of-whole analyses to determine **(B)** composition of cell types on vessels at each age, and **(C)** the cell type found at the greatest depth on individual vessels at each age. **(D-E)** Graphs depicting the (**D)** maximum depth of the furthest-migrated PVF and PVM on vessels measured from the meninges at ages P6, P7, P8 and P9/10 and **(E)** percent coverage of vessels by each cell type at each age. For B-C, the means of each biological replicate (n = 3 for each age) are represented by large colored shapes (cyan = PVFs, yellow = PVMs) and corresponding technical replicates (n = 10-50 vessels per animal) in small gray matching shapes, line and error bars show mean and SD. One-way ANOVA with multiple comparisons of PVF and PVM groups at each age reveal no statistical significance in depth or coverage measurements between cell types from P6 to P9/10, but revealed statistical significance between PVM depth and coverage across timepoints (**C)** P6 vs. P9/10, p = 0.0097; P8 vs. P9/10, p = 0.0354; **D)** P6 vs. P9/10 p = 0.0009; P8 vs. P9/10 = 0.0353; p < 0.05 *, p < 0.01 **, p < 0.001 ***).

Progression of PVM coverage of vessels from P6 to P10 followed a strikingly similar pattern as PVFs on the same vessel (**Fig. 4A**, arrowheads). To quantify the dynamics of coverage of PVFs and PVMs, we analyzed the percent coverage of PVFs and PVMs on individual vessels at P6, P7, P8, and P9/10, and found similar trends in the two cell types (**Fig. 4E**). Like PVFs, PVMs significantly increased coverage of vessels from P6 to P9/10 (p = 0.0009, p < 0.001 ***), with a mean of 93% PVM coverage by P9/10 (**Fig. 4E**). PVFs had a slightly higher average percent coverage at each age, however there were no statistical differences between the percent coverage of PVFs and PVMs on vessels at each timepoint (**Fig. 4E**). We also compared density of PVFs and PVMs on vessels within the covered regions and found that PVFs and PVMs had approximately the same density across timepoints, suggesting they are present in a roughly 1-to-1 ratio during postnatal development (**Fig. S3B**). Similar to PVFs, PVMs do not change in their density on vessels from P6-P9/10 yet they increase in percent coverage over these timepoints, meaning PVMs increase in number to cover a greater vessel length (**Fig. S3B**). This aligns with previous work showing clonal expansion of PVMs along vessels from P1 to P14, suggesting they proliferate locally to cover vessels during this developmental window (Masuda et al., 2022). Taken together, these data show that two different perivascular cell types, macrophages, and fibroblasts, develop in perivascular spaces largely in tandem in the postnatal brain.

### Lineage tracing reveals clonal domain patterns of brain PVFs on cortical blood vessels

PVFs located on vessels in the brain after P5 appear to emerge from a population of PVFs positioned along the meningeal vasculature during embryonic development and prior to P5 (**Fig. 2-3**). We next wanted to understand the clonal lineage of cerebral PVFs and determine if there were specific patterns to PVF expansion on vessels. First, we predicted that meningeal PVFs directly produce brain PVFs on nearby penetrating vessels. To test this, we used the *Col1a2-CreERT* line along with the *R26R-Confetti(Brainbow)-flox* reporter line, which labels recombined cells stochastically with one of four fluorescent proteins. We also determined a tamoxifen dosage in which the *Col1a2-CreERT* would induce sparse recombination in CNS fibroblasts, such that we observed only 2-3 spatially disparate clones within a 5x field of view in each sample. This system allowed us to identify groups of cells, or clones, that arise from a single, recombined cell. We injected *Col1a2-CreERT;Brainbow-flox* pups at P1 with a sparse labeling dose of tamoxifen, and collected brains at P14, a timepoint in which all cerebral vessels are covered by PVFs (**Fig. 5A**). In P14 brains, we observed a population of recombined cells exclusively localized in the meninges, such as the *RFP+* cells in the representative image shown (**Fig. 5A**, meningeal clones, arrows). These labeled cells are likely recombined non-perivascular meningeal fibroblasts and are never observed with corresponding clones on adjacent cerebral vessels. We also observed recombined brain PVFs covering vessels such as the *YFP+* PVFs shown (**Fig. 5A**, perivascular clones, arrowheads) that share a clonal origin with a recombined cell on a lectin+ vessel in the leptomeninges located directly above the cerebral vessel (**Fig. 5A**, asterisk). Taken together, our lineage tracing using both the *tdTomato-flox* and *Brainbow-flox* reporter lines show that brain PVFs developmentally originate from meningeal PVFs located proximal to the junction between the meningeal vessel and corresponding cerebral penetrating vessel.

**Figure 5:**
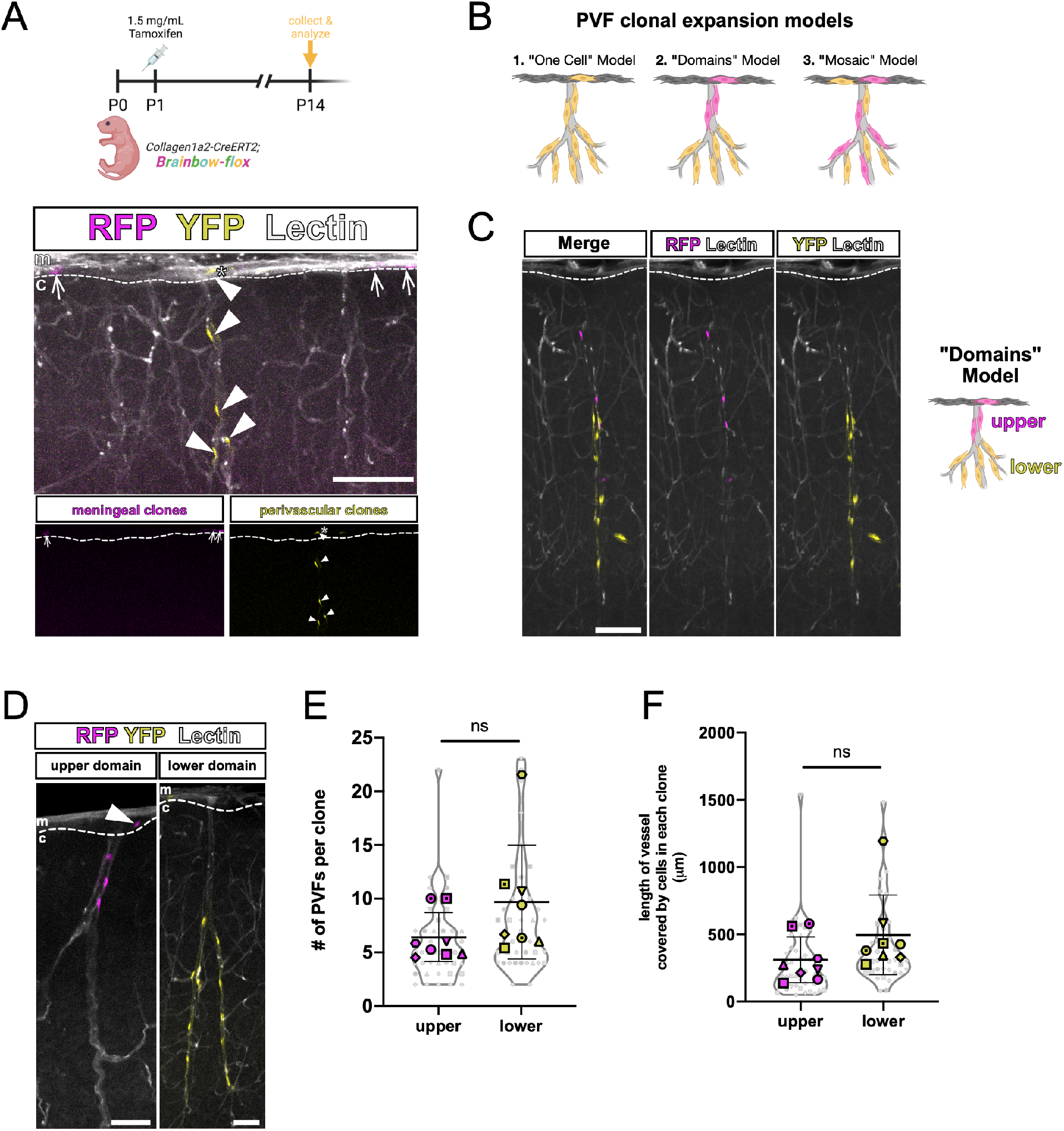
Parenchymal PVFs originate in the meninges and undergo regional clonal expansion on vessels in the brain. **(A-D)** Lineage tracing of fibroblasts in CUBIC-cleared *Col1a2-CreERT2;Brainbow-flox* brains. High-magnification images of vessels (Lectin, white) and sparsely-labeled fibroblast clones recombined at P1 and analyzed at P14 (RFP and YFP). **(A)** Arrows mark RFP+ meningeal fibroblast clones, arrowheads mark YFP+ brain PVF clones containing a meningeal-localized PVF (asterisk), insets show RFP and YFP clones only. Meninges, **m**; cortex, **c**. Scale bar 100µm. **(B)** Cartoon depiction of potential models of PVF coverage of parenchymal vessels. **1.** One-cell Model: a single meningeal clone gives rise to all cells on an individual vessel. **2.** Domains model: multiple clones account for coverage of individual vessels, and clones have regionalized domains. **3.** Mosaic model: clones stochastically cover vessels in a mosaic fashion. **(C)** Image shows individual vessel containing two PVF clones in respective upper (RFP, magenta) and lower (YFP, yellow) domains. Meninges, **m**; cortex, **c**. Scale bar 100µm. **(D)** Representative images of upper (RFP, magenta) and lower (YFP, yellow) domain clones. Arrowhead marks a *RFP+* meningeal PVF belonging to the ‘upper’ domain clone. Meninges, **m**; cortex, **c**. Scale bars 100µm. **(E-F)** Graphs depicting **(E)** number of PVFs in each clone and **(F)** length of vessel covered by a single clone, categorized by domain type and with the mean of each biological replicate (n = 8 animals) represented by large colored shapes and corresponding technical replicates (n = 5 – 22 clones per animal) in small gray matching shapes, line and error bars show mean and SD. Unpaired t-tests between each group revealed no significant differences between domain types.

In the *Col1a2-CreERT;Brainbow-flox* brains, we observed brain PVFs from the same clonal origin such as the *YFP+* clone shown in **Fig. 5A**, however we noted that the labeled clone never covered the entire length of the penetrating vessel. We know from observing the *Col1a1-GFP* reporter at P14 and *Col1a2-CreERT;tdTomato-flox* lineage tracing experiment that all penetrating cerebral vessels contain meninges-derived PVFs at this timepoint and are nearly fully covered (**Fig. 1A & C**, **Fig. 2**). Thus, there must be unlabeled clone(s) covering the rest of the vessels we observed. This made us speculate on whether there were certain patterns to how PVFs tiled on to cerebral vessels, and we first generated models to explain how this might occur (**Fig. 5B**). The first “One-Cell Model” dictates that all brain PVFs on a singular vessel share a clonal origin with one meningeal PVF (**Fig. 5B**, 1), though we can immediately rule out this model because of the empty spaces containing unlabeled cells that we noticed on vessels. Thus, two or more PVF clones must occupy an individual vessel to achieve full coverage. The “Domains Model” suggests that PVF clones cover vessels in a patchwork fashion, occupying non-overlapping domains (**Fig. 5B**, 2), while in the “Mosaic Model” PVF clones cover vessels in stochastic patterns (**Fig. 5B**, 3). In the brains from *Col1a2-CreERT2;Brainbow-flox* pups injected with tamoxifen at P1 and collected at P14, we find that PVF clones occupy domains as described by the “Domains Model” (**Fig. 5C-D**). Here, we can categorize recombined PVFs of a single clonal origin into ‘upper’ or ‘lower’ clone types, based on their location either on the upper or lower part of the vessel (**Fig. 5C-D**). In rare instances, we observe vessels with multiple PVF clones that are spatially separate (**Fig. 5C**). ‘Upper’ domain clones are always observed with a meningeal clone directly adjacent to the vessel, while ‘lower’ domain clones often do not have an observable corresponding meningeal clone within the imaging area (**Fig. 5D**). We speculate that this reflects initiating PVFs that originate on vessels in the meninges, but between P5 and P14 migrate deeper on the vessel and proliferate to give rise to other PVFs locally. To further characterize ‘upper’ and ‘lower’ PVF clones, we counted the number of cells per clone and the length of vessel covered by each clone (**Fig. 5E-F**). ‘Lower’ clones tend to contain more cells (mean = 10 PVFs) compared to ‘upper’ clones (mean = 6 PVFs), and ‘lower’ clones tend to cover more vessel length (mean = 497.8µm) compared to ‘upper’ clones (mean = 312.7µm), though neither of these differences are statistically significant (**Fig. 5E-F**). To summarize, PVFs located on vessels in the cortex developmentally arise from PVFs on meningeal vessels near junctions with penetrating vessels, and multiple meningeal PVFs are required to give rise to all PVFs that cover each vessel. Brain PVFs of the same clonal origin cluster on vessels in distinct non-overlapping domains, and domain location potentially reflects differences in the timing of emergence and migration of individual PVFs.

### PVFs migrate and proliferate to achieve full coverage of cerebral vessels

We have shown that brain PVFs originate from the meninges and undergo clonal expansion to cover vessels postnatally (**Fig. 5**). Inherent to this model of PVF coverage of vessels is that PVFs must migrate and undergo cell division, thus we next investigated migration and proliferation of PVFs to understand how these cellular mechanisms fit in to their developmental progression. We first used *in vivo* two-photon imaging of *Col1a1-GFP+* mice at P8 to look for evidence of PVF migration during postnatal development (**Fig. 6A**). We selected P8 because it falls within the temporal range (P7-P10) where we see the greatest increases in cerebral depth and percent coverage of vessels (**Fig. 1B-C**). A thinned-skull cranial window was created in *Col1a1-GFP+* pups at P7, then pups were imaged twice on P8 at intervals 6 hours apart. Intravenous fluorescent dye (2MDa Alexa-680) was used to visualize blood vessels (**Fig. 6A-B**). From large overview scans of the meningeal surface, we were able to identify and image penetrating vessels containing PVFs and annotate the locations of the PVFs along penetrating vessels at each timepoint (**Fig. 6B-B’’**). Using this information, we calculated the displacement of individual PVFs on vessels and found that most PVFs displace positively, meaning they move downward on vessels into the cortex (**Fig. 6C**). This can be compared to measurements of PVFs in adult mouse brains obtained using similar *in vivo* imaging, adult PVFs never displace more than 5-10µm when imaged repeatedly over weeks (Bonney et al., 2022). During our analysis, we noted that PVFs located deeper on vessels seemed to make more dynamic movements compared to PVFs located closer to the meningeal surface, which seemed to make more minor movements. We classified the deeper, more dynamic PVFs as “Leading PVFs” and the more superficial PVFs as “Trailing PVFs” (**Fig. 6B’-B’’**, asterisks). When we binned displacement values for PVFs based on whether they are Leading or Trailing, we found that Leading PVFs have a significantly higher mean displacement of 11.2µm compared to Trailing PVFs with a mean displacement of 1.5µm over the 6-hour interval (p = 0.0003, p < 0.001 ***) (**Fig. 6D**). We also calculated total displacement over time of Leading and Trailing populations and found that Leading PVFs traveled greater total distances from t = 0hr to t = 6hr compared to Trailing PVFs (p = 0.0159, p < 0.05 *) (**Fig. 6E**). Collectively this data support that PVFs migrate to achieve coverage of penetrating cerebral vessels, and there are differences in migratory dynamics based on the position of a PVF on the vessel (trailing vs leading).

**Figure 6:**
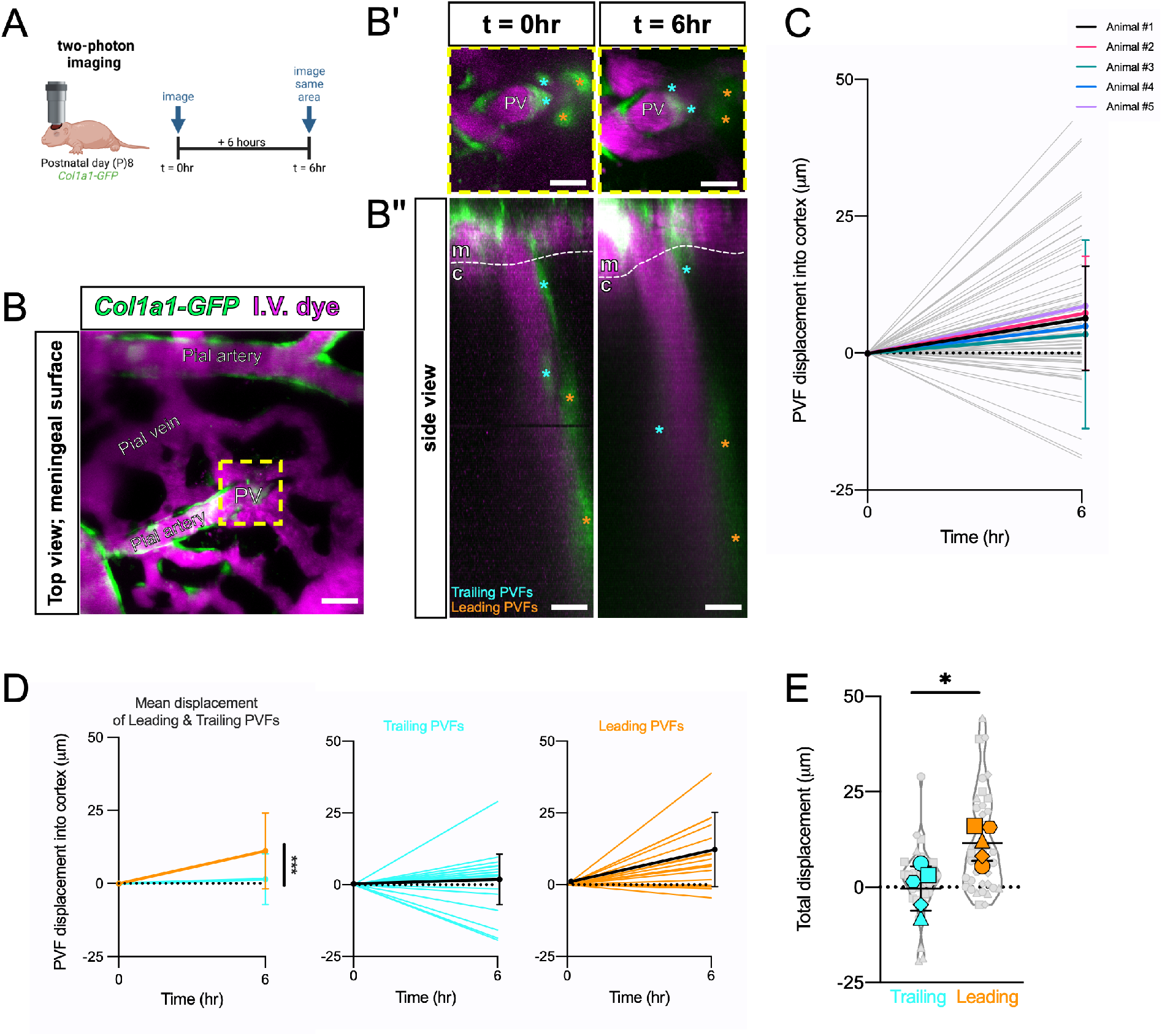
*Col1a1-GFP+* PVFs migrate along vessels in P8 mice *in vivo*. **(A)** Schematic of imaging paradigm; *Col1a1-GFP+* P8 mice installed with cranial windows were imaged using two-photon microscopy at two timepoints 6 hours apart (t = 0hr and t = 6hr). **(B)** Overview image of the meningeal surface showing fibroblasts (GFP, green) and vessels (IV dye, magenta). Penetrating Vessel, **PV**. Insets show high-magnification zoom **(B’)** and side view projection of the zy-plane **(B’’)** of area in yellow box, with individual leading PVFs (orange asterisks) and trailing PVFs (cyan asterisks) annotated at t = 0hr and t = 6hr. Meninges, **m**; cortex, **c**. Scale bars 25µm. **(C-E)** Graphs showing displacement of individual PVFs on vessels into the cortex between t = 0h and t = 6hr, lines and error bars show mean and SD. **(C)** Summary of all PVFs measured (gray lines, n = 8 – 22 PVFs per animal) and means for each biological replicate (colored lines, n = 5 animals). Two-way ANOVA with multiple comparisons shows no significant differences between animal replicates over time. **(D)** Displacement data split by leading PVFs (orange lines, black line shows mean) and trailing PVFs (cyan lines, black line shows mean). Two-way ANOVA analysis of trailing and leading PVFs revealed a statistically significant difference in displacement between trailing and leading populations over time (p = 0.0003, p < 0.001 ***). **(E)** Graph showing total calculated displacement (µm) from t = 0hr to t = 6hr of leading (orange) and trailing (cyan) PVF populations in *Col1a1-GFP+* mice at P8. The means of each biological replicate (n = 5 animals) are represented by large colored shapes and corresponding technical replicates (n = 8 – 22 PVFs per animal) in small gray matching shapes, line and error bars show mean and SD. Unpaired t-test analysis reveals a statistically significant difference in the mean displacement over time of these two groups (Leading vs. Trailing, p = 0.0159, p < 0.05 *).

Migration partially accounts for the ability for PVFs to cover vessels, however we also observe expansion of PVF clones in our lineage tracing experiments (**Fig. 5**), and PVFs maintain an equal density while increasing in coverage (**Fig. S1B**), thus, PVFs must undergo cell division during their development. To determine the timing of PVF proliferation during development, we performed 2-hour EdU pulse-chase assays in *Col1a1-GFP+* mice at embryonic and postnatal timepoints (**Fig. 7**). We first quantified proliferation of leptomeningeal PVFs using leptomeningeal flat-mounts from *Col1a1-GFP+* mice injected 2 hours prior to collection with EdU at ages E16, P0, and P2 and counted the number of GFP+/EdU+ PVFs on leptomeningeal vessels (**Fig. 7A-B**, A inset and arrowheads). The average percent of GFP+/EdU+ PVFs between E16 and P2 fell between a range of 48% and 57% and did not significantly change across timepoints (**Fig. 7B**). Possibly, meningeal PVFs ‘prime’ themselves for emergence onto cerebral vessels by proliferating during embryonic and early postnatal timepoints to maintain high density and readiness to migrate into perivascular spaces. We then performed the same experiment on *Col1a1-GFP+* pups at P7, P9, P11, and P14 and collected brains for CUBIC-clearing and counting of GFP+/EdU+ PVFs on penetrating vessels (**Fig. 7C-D**). We found a significantly higher percentage of GFP+/EdU+ PVFs on vessels in the brain at P7 compared to later timepoints suggesting P7 is a proliferative period for PVFs (**Fig. 7D**). In general, brain PVFs were much less proliferative between P7 and P14 than meningeal PVFs at earlier timepoints (**Fig. 7B**). These data suggest meningeal PVFs increase in number to ‘prepare’ for migration, then once on cerebral vessels undergo low levels of proliferation on vessels over the span of a week to achieve vessel coverage. Interestingly, the P7 proliferative window for PVFs is just before when we observe *in vivo* migration of PVFs, thus we speculate that brain PVFs may migrate and proliferate concurrently. Collectively, this data supports a mechanism in which PVFs enter the brain via penetrating vessels at P5, migrate and proliferate locally on vessels between P7 and P10, then further refine their positioning and extend on to branches between P10 and P14 (**Fig. 8**).

**Figure 7:**
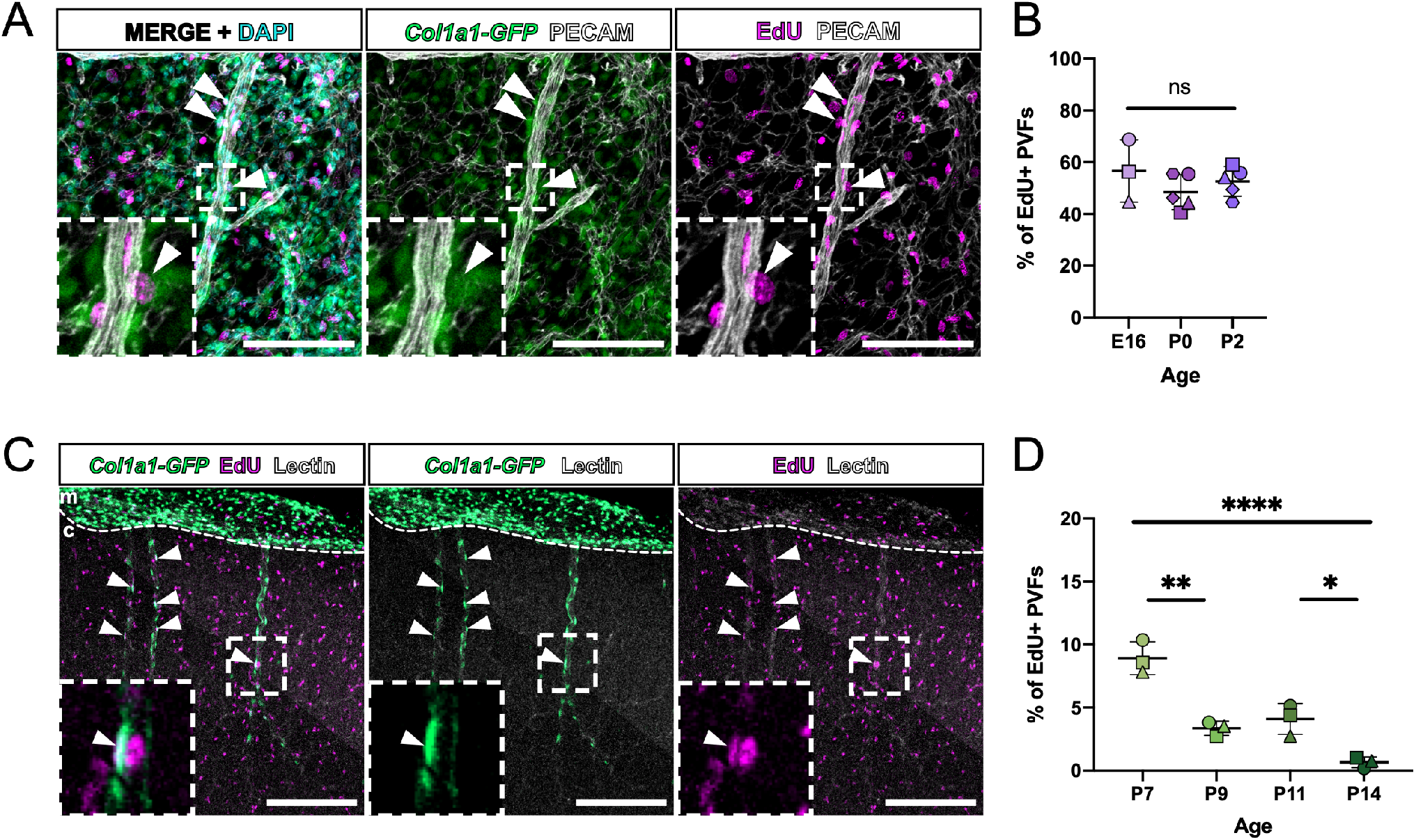
*Col1a1-GFP+* PVFs proliferate along vessels in the meninges and brain during embryonic and postnatal development. **(A)** Flat-mount preparation of P5 meninges showing fibroblasts (GFP, green), vasculature (PECAM, white), EdU+ nuclei (magenta), and nuclei (DAPI, cyan). Inset and arrowheads mark meningeal PVFs that are EdU+. Scale bars 100µm. **(B)** Graph showing the percentage of EdU+/GFP+ PVFs at ages E16, P0, and P2. Line and error bars show mean and SD. One-way ANOVA with multiple comparisons detects no statistically significant differences in the percentage of EdU+/GFP+ PVFs between E16, P0, and P2. Experiments were performed with biological replicates of n = 3 – 5 for each age, and for each animal replicate n = 96 – 1230 PVFs were counted. **(C)** High-magnification images of CUBIC-cleared *Col1a1-GFP* brains at P8 showing fibroblasts (GFP, green), vessels (Lectin, white), and EdU+ nuclei (magenta). Insets and arrowheads mark PVFs that are EdU+. Scale bars 200µm. **(D)** Graph showing the percentage of EdU+/GFP+ brain PVFs at ages P7, P9, P11, and P14. One-way ANOVA with multiple comparisons revealed statistically significant differences between timepoints (P7 vs. P9, p = 0.0013; P7 v.s. P14, p < 0.0001; P11 v.s. P14, p = 0.0103; p < 0.05 *, p < 0.01 **, p < 0.001 ***, p < 0.0001 ****). Experiments were performed with biological replicates of n = 3 for each age, and for each animal replicate n = 70 – 734 PVFs were counted.

**Figure 8:**
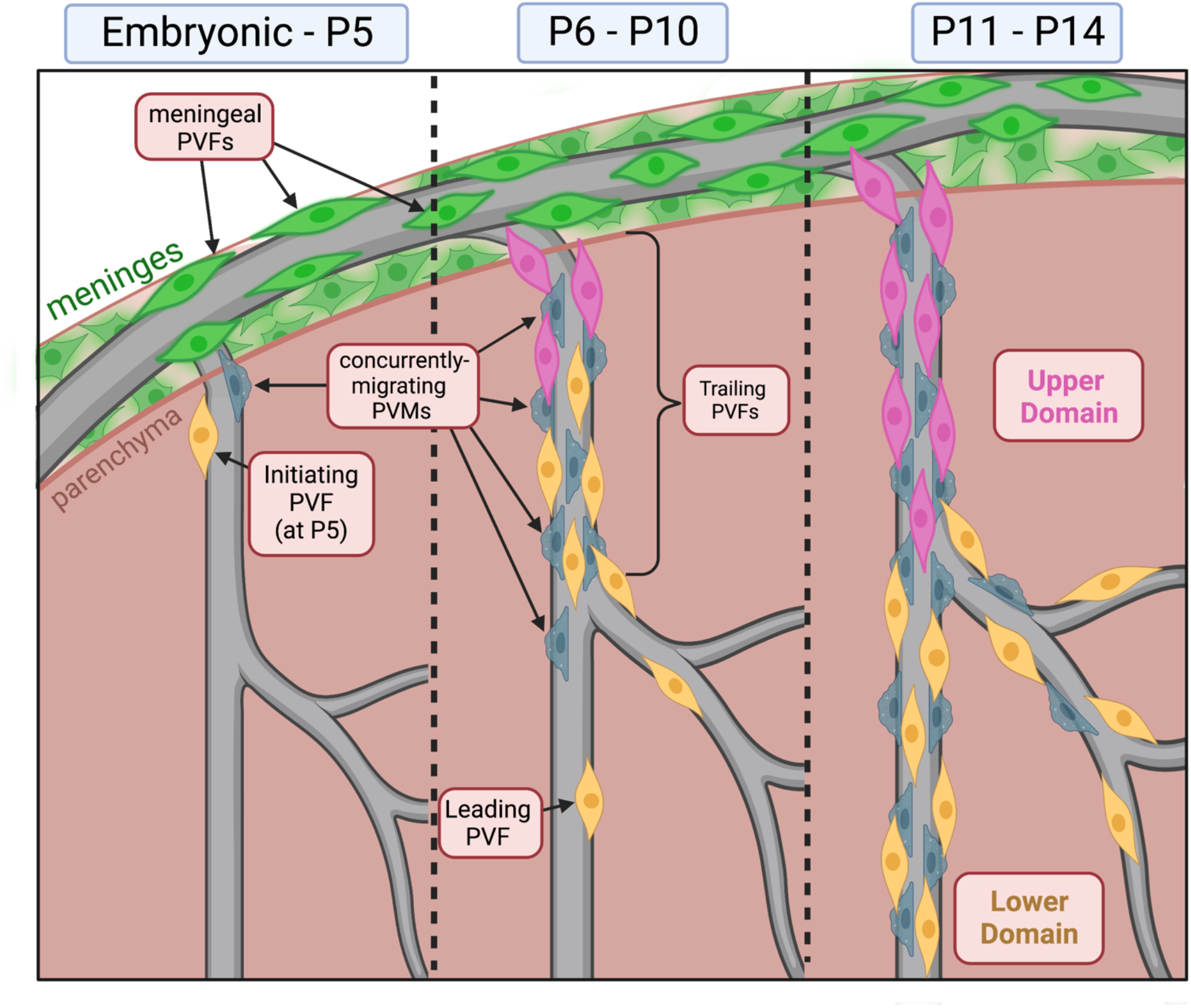
Summary of brain perivascular fibroblast development. PVF development occurs in a window between P5 and P14 in the postnatal mouse brain. This process occurs concurrently with PVM development on vessels. Prior to P5, PVFs are absent from the brain and are located exclusively on meningeal vessels. At P5, initiating PVFs begin migrating on vessels. Between P6 and P10, PVFs migrate and proliferate on vessels. At P8, we find distinct subsets of Leading and Following PVFs based on their average displacement and velocity. Between P11 and P14, PVFs continue to migrate and proliferate to fully cover the vessel. PVF coverage can be separated in to “upper” and “lower” domains based on clonal origin.

## DISCUSSION

Here, we define PVF development between P5 and P14 in the cerebral cortex and identify the cellular mechanisms by which PVFs cover vessels; from this, we can build a comprehensive timeline of PVF development during postnatal development (**Fig. 8**). At P5, PVFs are observed initiating coverage of penetrating vessels, between P6 and 10 they migrate and proliferate to cover the main branches of vessels, and by P14 they have fully covered the main branches and have extended on to secondary branches (**Fig. 8**). We also show that PVMs follow a similar developmental trajectory (**Fig. 8**). Analysis of leptomeningeal flat-mounts combined with lineage tracing using *Col1a2-CreERT2* mice supports a meningeal origin for brain PVFs. We also find that PVFs expand clonally on vessels and occupy non-overlapping domains, and the locations of these domains may correlate with differential cell migratory dynamics and timing of emergence on the vessel as shown by *in vivo* observation (**Fig. 8**). Altogether, this work sheds light on a poorly understood developmental process and may provide us with clues about the functions of PVFs within the neurovascular unit.

The entirely postnatal timing of PVF appearance on vessels in the cerebral cortex is particularly striking, given that the ‘scaffold’ of the neurovascular unit (endothelium, pericytes/vSMCs) upon part of which PVFs migrate is established days earlier, during prenatal development (Paredes et al., 2018). PVF coverage of cerebral vessels occurs in parallel with appearance of PVMs, and we show both cell types have nearly identical progressive coverage of cerebral vessels. Further, PVFs and CD206+ leptomeningeal macrophages, which give rise to CD206+ PVFs (Masuda et al., 2022), are in high abundance in the leptomeninges for days prior to their appearance on cerebral vessels. What events are occurring in the cerebral vasculature that could account for the timing of PVF and PVM moving onto cerebral vessels? Possibly, it could relate to the postnatal appearance of perivascular spaces, a gap that forms around large diameter vessels between the vSMC-covered endothelium and astrocyte endfeet that contains interstitial fluid and is continuous with the subarachnoid space of the meninges (Wardlaw et al., 2020; Zhang et al., 1990). Perivascular spaces play a critical role in waste clearance and fluid homeostasis as part of the CNS “glymphatic” system (Plog and Nedergaard, 2018; Smith et al., 2017), and studies in humans and rodents indicate PVS function is necessary for maintaining brain health (Francis et al., 2019; Iliff et al., 2012). Interestingly, astrocyte endfeet polarization of water channel Aquaporin 4, and the presence of interstitial fluid in perivascular spaces, a key characteristic of the glymphatic system, emerges between P7 through P14 in the mouse cortex (Munk et al., 2019). Recent work on PVM development in mice reported that perivascular spaces near and distal from the pial surface widen between P4 and P9 (Masuda et al., 2022). These observations support postnatal appearance of perivascular spaces, thus creating a space for PVF and PVM to migrate into on the vessel surface. Additionally, extensive expansion of the cerebral capillary network occurs between P8 and P12 (Coelho-Santos et al., 2021a). Possibly, ongoing cerebrovascular development to support rapid increases in metabolic demand of the postnatal brain may create permissive signals to attract PVFs onto the cerebral vasculature. Further work is required to detail how the development of PVFs corresponds with emergence of perivascular spaces and the postnatal maturation of the neurovascular unit.

Our lineage tracing data and *in vivo* imaging of developing PVFs identified specific spatiotemporal patterns of PVF emergence. ‘Lower’ domain PVF clones, seen covering the lower branches of vessels, may correspond to ‘leading’ PVFs seen displacing further and migrating faster to reach vessel termini and expand locally, whereas ‘upper’ PVF clones localized more proximal to the meningeal surface may correspond to ‘trailing’ PVFs that have a lower average displacement. We speculate that the distinction between these two PVF sub-types relates to their proximity to junctions between meningeal and penetrating vessels prior to P5; meningeal PVFs close to these junctions will be the first to initiate coverage at P5, and remaining PVFs on the meningeal vessel will follow behind. Whether ‘leading’ and ‘trailing’ PVF populations have distinct molecular and functional characteristics is unknown. In models of neural crest and cancer cell collective migration, leader cell populations have distinct morphology and cytoskeleton organization (Qin et al., 2021), and in trunk neural crest, ablating leader cells results in failure of follower cell migration (Richardson et al., 2016). Potentially, ‘leading’ and ‘trailing’ PVFs have differential propensities to respond to molecular signals, explaining their differential migratory capacity.

PVF response to CNS injury, which includes both enhanced ECM and signaling molecule production, provides some insight into the functional significance of postnatal PVF development. In spinal cord injury and stroke, PVF numbers expand dramatically and deposit ECM proteins that form the fibrotic scar (Fernández-Klett et al., 2013; Kelly et al., 2016; Soderblom et al., 2013). In healthy tissues, fibroblasts in many organs function in producing ECM, thus it is possible that PVFs may function in secreting, maintaining, and modifying ECM during development and homeostasis. This potential function of PVFs may also be tied to the development of PVMs. During postnatal development, brain PVMs rely on integrin-dependent mechanisms to populate parenchymal vessels (Masuda et al., 2022). Potentially, PVFs function to contribute to local ECM for integrin-dependent migration of PVMs. This may also explain why PVFs were more frequently detected ahead of PVMs, possibly laying down a permissive environment for PVM migration. PVFs may also function in producing molecular signals required for development and maintenance of the PVS and neurovasculature. Our lab has shown previously that in a mouse model of stroke, in addition to their fibrotic role, PVFs producing retinoic acid (RA) in the site of injury (Kelly et al., 2016). Developmental factors produced by meningeal fibroblasts, in particular RA, are required for cortical development (Siegenthaler et al., 2009) and cerebrovascular development (Bonney et al., 2016; Mishra et al., 2016). We found previously that adult brain PVFs express some of the components of the RA synthesis pathway (Raldh1/Raldh2) (Kelly et al., 2016), and we show here that meningeal-located PVFs express Raldh2. It is possible that PVFs are a local source for RA in the brain during postnatal development, and their heightened production of RA during injury reflects a re-activation of developmental pathways.

Our study provides an important framework for the cellular mechanisms driving PVF development in the mouse brain. A major outstanding question is: what are the molecular mechanisms controlling PVF development? We highlight integrins/ECM modification and RA signaling as potential functions for PVFs; it is possible that PVFs require these mechanisms for their own development as well. Us and others have previously shown that PVFs express both PDGF receptors α and β (Fernández-Klett et al., 2013; Kelly et al., 2016; Riew et al., 2020; Vanlandewijck et al., 2018). PDGF-B ligand produced by the endothelium is required for recruitment of pericytes to the brain vasculature during embryonic development (Lindahl et al., 1997); it is possible that PVFs rely on a similar mechanism for their progression on the vasculature during postnatal development. Additionally, interstitial fluid flow within perivascular spaces may play a role by providing the molecular cues or mechanical stimulus required for PVF migration and proliferation on vessels. Given their positioning within perivascular spaces, it is very likely PVFs receive their developmental cues via cell-cell signaling with other proximal neurovascular cell types. Future studies aimed at understanding the specific signals required for PVF development, and how PVFs fit into the overall development of PVSs and the neurovascular unit are now possible using our model of the timing and cellular mechanisms of PVF development.

## METHODS

### Animals

Mice used for experiments were housed in specific pathogen-free facilities approved by AALAC and procedures were performed in accordance with animal protocols approved by the IACUC at The University of Colorado, Anschutz Medical Campus and Seattle Children’s Research Institute. Mouse lines used in this study include: 1) *Col1a1-GFP*: Tg(Col1a1-EGFP)#Dab (MGI: 4458034 (Yata et al., 2003)), 2) *Col1a2-CreERT2*:*Tg*(Col1a2-cre/ERT,-ALPP)7Cpd (Jackson Strain #:029567, RRID:IMSR_JAX:029567), 3) *Ai14(tdTomato)-flox*: B6.Cg-*Gt(ROSA)26Sor^tm14(CAG-tdTomato)Hze^*/J (Jackson Strain #:007914, RRID:IMSR_JAX:007914), 4) *Brainbow-flox*: *Gt(ROSA)26Sor^tm1(CAG-Brainbow2.1)Cle^* (Jackson Strain #:013731, RRID:IMSR_JAX:013731). For flat-mount and cleared tissue analyses, adult mice were crossed to generate embryonic (E12 – E16) or postnatal litters collected at ages E14, E16, P0, P2, P3, P5, P6, P7, P8, P9, P10, P11, and P14. For lineage tracing experiments, P1 mice were injected into the milk sac with 50 µL of 1 mg/mL (*Ai14-flox*) or 1.5 mg/mL (*Brainbow-flox*) of tamoxifen (Sigma, Cat#: T5648) in corn oil (Sigma, Cat#: C8267) to achieve Cre-activated recombination. For two-photon imaging studies on P8 *Col1a1-GFP* pups, we followed our previous methods that are comprehensively described in *Coelho-Santos et al. 2021* and *Coelho-Santos et al. 2022* (Coelho-Santos et al., 2021a, 2022). For EdU-based proliferation assays, we intraperitoneally injected 2.5 mg/mL EdU (Thermo Fisher, Cat#: NC1495577) into pregnant mice for embryonic collections (150 µL) or postnatal mice (50 µL) 2 hours prior to collection.

### Tissue collection and immunohistochemistry

All procedures and reagents used for collecting and processing meningeal whole mounts and cleared brain slices for immunohistochemistry, EdU detection, and imaging are described at length in our methods paper (Jones et al., 2022). The following primary antibodies were used in meningeal whole mount and cleared tissue staining: rabbit anti-Desmin (Cell Signaling, Cat#: D93F5), rat anti-CD31/PECAM (BD Pharmigen, Cat#: 561814), rabbit anti-S100A6 (Novus, Cat#: NBP1-89388), rabbit anti-RALDH2 (Sigma, Cat#: HPA010022), and goat anti-CD206 (R&D Systems, Cat#: AF2535). Appropriate species-specific Alexa-Fluor secondary antibodies (Invitrogen) were used in all cases. In cleared tissue, vasculature was labeled by DyLight Tomato Lectin 649 (Vector Laboratories, Cat# DL-1178-1). EdU detection was performed with the Click-iT Plus EdU Alexa Fluor 647 Imaging Kit (Thermo Fisher, Cat#: C10640). Images were obtained using a Zeiss 900 LSM and Zen Blue software and processed in FIJI.

### Image acquisition and quantification

Images were obtained using a Zeiss 900 LSM and Zen Blue software. All processing and image quantification was performed in FIJI. For analysis of cell depth, percent coverage, and cell density in **Figures 1 & 4**, individual vessels were selected for analysis by observing the labeled vessel channel only (lectin, far red) and annotating penetrating vessels in which the entire length of vessel could be seen from the meninges to the terminus. For cell depth analysis, the distance from the surface of the meninges to the soma of the furthest-migrated cell on the vessel was measured using the segmented line tool in FIJI. For percent coverage measurements, the depth measurement (PVF or PVM “coverage” length) was divided by the total length of the vessel. For density measurements, the coverage length was divided by the number of cells found within the covered length of the vessel then multiplied by 100 µm to normalize measurements to a given length. Similar analyses were conducted for density measurements of meningeal PVFs (**Fig. 3D**); PVFs characterized by *Col1a1-GFP* labeling and elongated morphology aligned to the vessel were counted over a measured length of large-diameter (> 8 µm) vessels, then multiplied by 100 µm to normalize measurements to a given length. For analyses of *Brainbow*-labeled PVF clones in **Figure 5E-F**, the number of PVF per labeled clone was counted to obtain “# PVFs per clone” and the length of vessel covered by each clone was measured using the segmented line tool in FIJI. For clones in which multiple branches of the vessel were covered, individual length measurements were taken and added together. For measurements of migratory dynamics of PVFs in **Figure 6**, detailed methods for *in vivo* image acquisition and calculation of PVF displacement from pial vessel branch points are described in *Bonney et al. 2022* (Bonney et al., 2022). Characterization of pial and penetrating vessel types were determined by branching patterns and vessel morphology as previously described (Coelho-Santos et al., 2021b). Leading PVFs were classified as the deepest two PVFs while the remaining, superficial PVFs were defined as trailing on the corresponding penetrating vessel. Further calculations of total displacement over time were conducted by summing displacement values for individual PVFs across timepoints. For analysis of cell proliferation in meninges flat-mounts and cleared brains in **Figure 6**, the number of EdU+ nuclei were counted and divided by the total number of GFP+ PVFs or meningeal cells counted across all images for each replicate, then multiplied by 100 to obtain a percentage.

### Statistical analysis

All statistical analyses were performed in GraphPad Prism (v.8.2.0). Normality tests were performed prior to statistical analyses where necessary. Respective statistical tests and replicate numbers are reported in figure legends. Where shown, lines and error bars represent mean and standard deviation.

## ACKNOWLEDGEMENTS

The authors would like to thank the members of the Siegenthaler Lab and members of HEJ’s thesis committee (Dr. Santos Franco, Dr. Katie Fantauzzo, Dr. Linda Barlow, Dr. Joe Brzezinski and Dr. Fabrice Dabertrand) for fruitful discussions and feedback that helped shape this project. This work was supported by funding from the NIH/NINDS (F31NS125875-01 to HEJ, R01 NS098273 to JAS; F32NS117649 to SKB), American Heart Association (Postdoctoral Fellowship to VCS), and NIH/NIGMS 1T32GM141742-01 support for the Developing Scholars summer research program for KAA and HEJ. All schematics found in the figures were made with BioRender.

## DECLARATION OF INTERESTS

The authors declare that they have no competing interests.

## Notes

### Competing Interest Statement

The authors have declared no competing interest.

