## Supplemental Figure 1-3 for "Meningeal origins and dynamics of perivascular fibroblast development on the mouse cerebral vasculature"

### SUPPLEMENTAL FIGURES

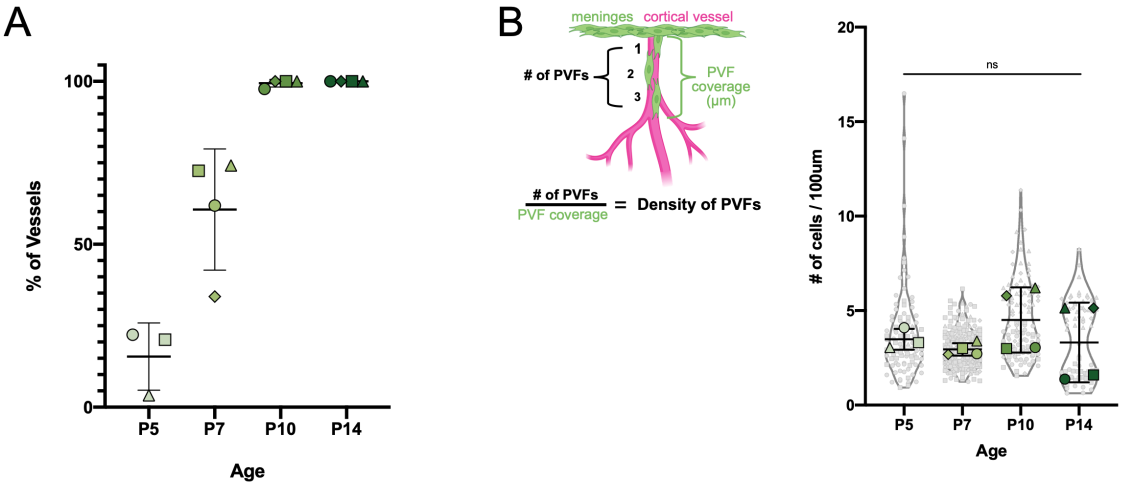


**Figure S1: Overall coverage of cerebral vessels by PVFs increases between P5 and P14, while density of PVFs on cerebral vessels does not change. (A)** Graph showing the percent of all vessels measured at ages P5, P7, P10, and P14 with any detectable PVF coverage; large colored shapes correspond to the mean for each individual biological replicate (n = 3 – 4 animals for each age), line and error bars show mean and SD. One-way ANOVA with multiple comparisons revealed a statistically significant increase between P5 and P7 (p = 0.0136) and P7 and P10 (p = 0.006). **(B)** Graph depicting density (number of cells per 100µm of vessel length) of PVFs at each age, with the mean for each biological replicate (n = 3 – 4 animals) represented by large colored shapes and corresponding technical replicates (n = 15-255 vessels per animal) in small gray matching shapes, line and error bars show mean and SD. One-way ANOVA with multiple comparisons revealed no significant changes across timepoints.

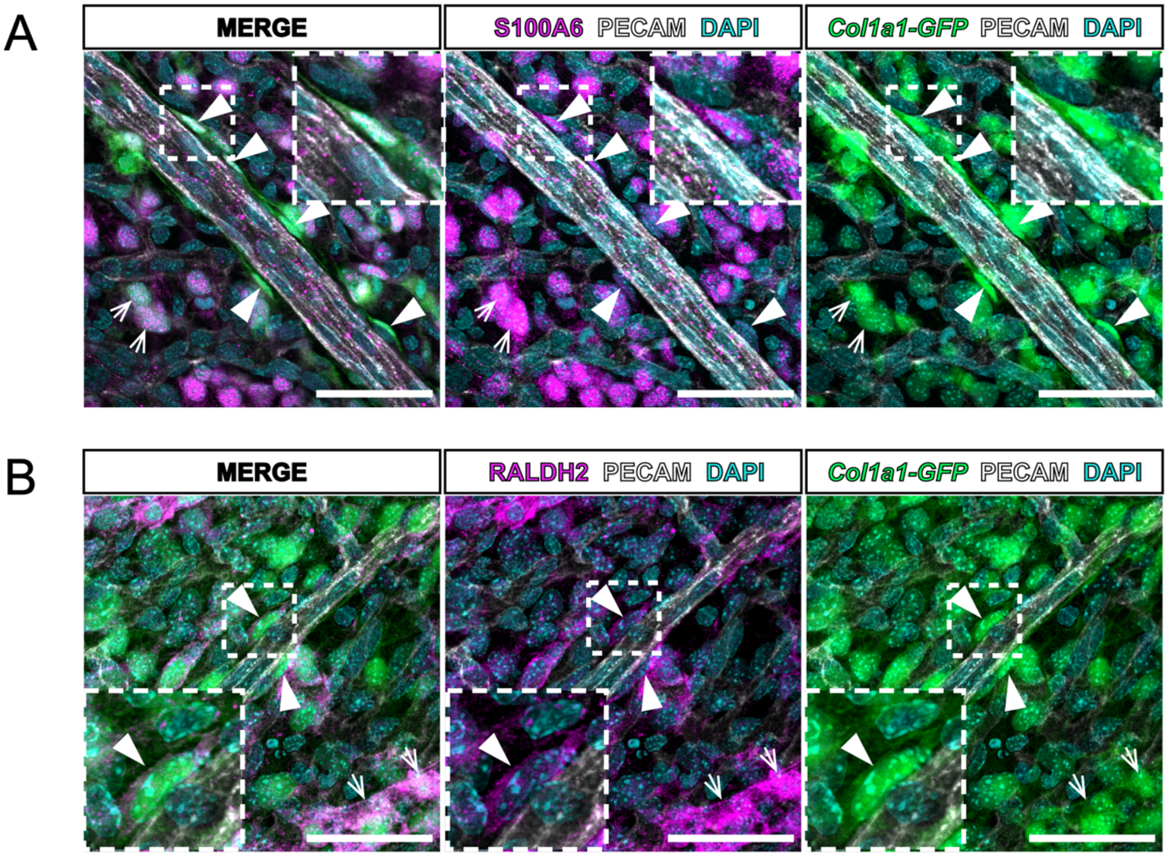


**Figure S2: PVFs in the cerebral leptomeninges express pia and arachnoid layer markers S100A6 and RALDH2.** Flat mount preparations of E15 leptomeninges showing fibroblasts (GFP, green), vasculature (PECAM, white), and **(A)** pia layer specific marker S100A6 (magenta), or **(B)** arachnoid layer marker RALDH2 (magenta). Non-perivascular meningeal fibroblasts expressing each of these markers are indicated with arrows. Meningeal PVFs express each of these markers, insets and arrowheads indicate **(A)** GFP+/S100A6+ PVFs or **(B)** GFP+/RALDH2+ PVFs. Scale bars 50μm.

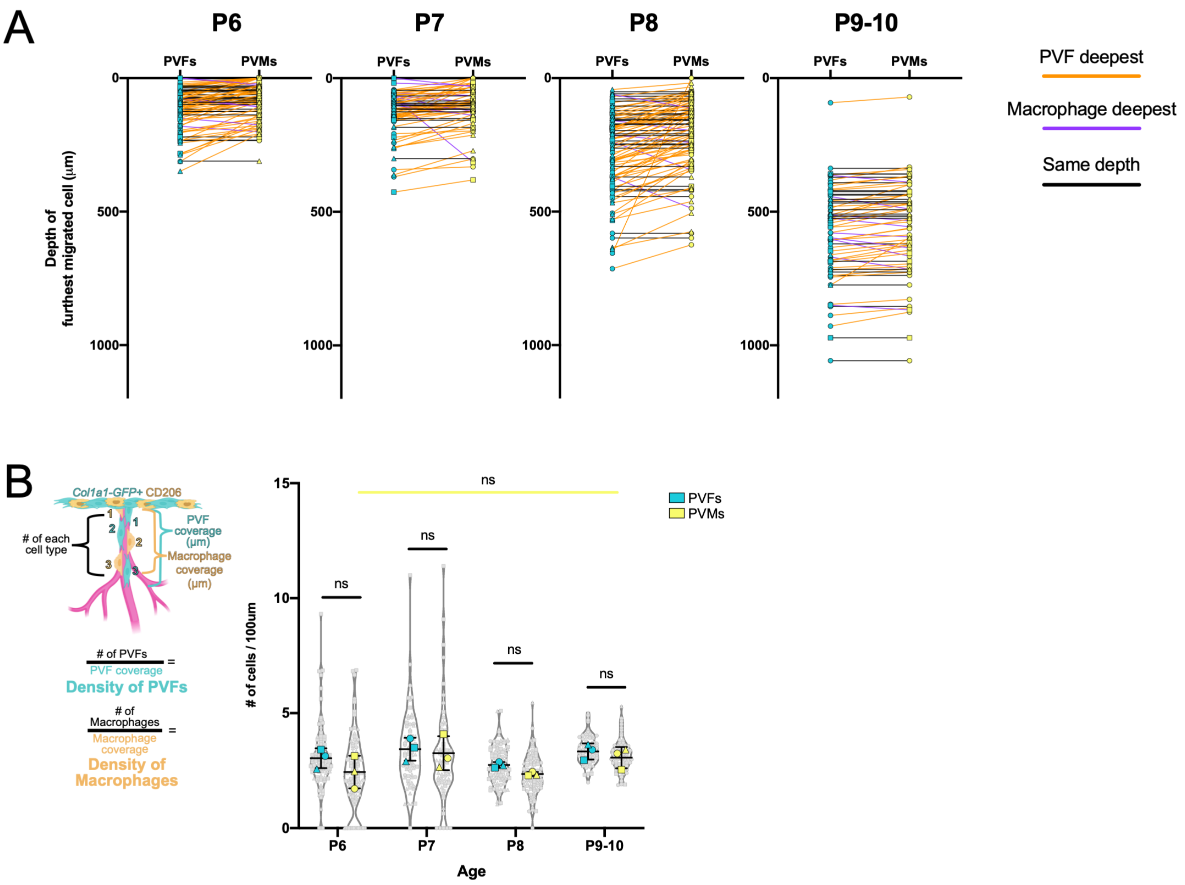


**Figure S3: Dynamics of PVF and PVM coverage during postnatal ages. (A)** Graphs showing the depth of PVFs and PVMs on individual vessels at P7 and P9/10. Each line-dot pair depicts and individual vessel, with the left dot (cyan) representing the maximum depth of PVFs and the right dot (yellow) representing the maximum depth of PVMs on that vessel. The color of the connecting line indicates which cell type is found deepest on each vessel (PVFs deepest, orange; PVMs deepest, purple; both at same depth, black). **(B)** Graph depicting density (number of cells per 100µm of vessel length) of PVFs and PVMs at each age, with the means of each biological replicate represented by large colored shapes and corresponding technical replicates in small gray matching shapes, line and error bars show mean and SD. One-way ANOVA with multiple comparisons of PVF and PVM groups at each age reveal no statistical difference in density between cell types at each age or for PVMs across timepoints. For analysis of A & B, each age used n = 3 biological replicates, with n = 10-50 technical replicates (vessels) analyzed per animal.
